# Virtual Touch: Sensing and Feeling with Ultrasound

**DOI:** 10.1101/2021.08.25.457633

**Authors:** Warren Moore, Adarsh Makdani, William Frier, Francis McGlone

## Abstract

The sense of touch codes the detection and properties of physical objects on the body via mechanoreceptors within the skin. Technological advancements, such as ultrasonic haptic devices, are now able to ‘touch without touching’, claiming this is virtual touch. An initial aim of the study was to investigate subjective intensity and pleasantness ratings of ultrasound stimulation and the influence of top-down factors using the Touch Experience and Attitudes Questionnaire (TEAQ). Self-reported intensity and pleasantness ratings were measured in response to ultrasound stimuli. A second aim was to record from individual low threshold mechanoreceptors using the technique of microneurography in an attempt to determine which mechanoreceptors are activated by ultrasound stimulation of the skin. The major findings here were that microneurography found SAI and SAII units did not respond to ultrasound stimuli; intensity and pleasantness ratings were significantly different between age groups. Ultrasound can produce a variety of sensations with varying intensity and pleasantness ratings. A limitation of the study was the unexpected force difference generated between modulations. These findings have implications for mid-air haptics, somatosensory affective research, and virtual reality. Future research should focus on microneurography investigation of FA fibre responses to ultrasound.

## Introduction

The sense of touch comprises four main sub-modalities; vibration/pressure, temperature, itch, and pain, which transmit neuroelectric signals to the brain via afferent nerves, collectively referred to as the somatosensory system (1). The term somatosensation originates from the Greek word *soma*, meaning body, and *sensation* meaning in this context, perception. In mammals the somatosensory system innervates the skin and muscles (2) but what we will focus on here is the cutaneous somatosensory system. In response to skin deformation a population of low threshold mechanoreceptors (LTM) transduce the mechanical energy into neuroelectric signals (spikes) in the afferent nerves, which then enter the spinal cord en route to the brain (3). Detecting the properties of physical objects through touch relies on the discriminative touch system, coded by fast conducting myelinated A-Beta fibres (~80 m/s and a diameter of ~10μm), which project to the somatosensory cortex (4, 5). Discriminative touch is largely found in the glabrous (palmar) skin of the hand and the perioral regions (lips/tongue). Cutting-edge technology is beginning to exploit current somatosensory knowledge to generate virtual touch, which describes replicating the sensation of an object touching the skin using focused ultrasound to stimulate LTMs (6). With the recent discovery of a system of c-low threshold unmyelinated mechanoreceptors (C-LTMs) the virtual replication of touch has entered a new domain (7) – C-LTMs are not found in glabrous skin and as their conduction velocities are ~1 m/sec they cannot serve a function in discriminative touch. The current theory on the functional role of C-LTMs are is that they code for the affective and rewarding properties of touch – a far more exciting target for the virtual world.

Afferent nerves are characterised by the degree of myelination, a method first established with electrophysiological studies of cats and monkeys (8, 9). The neurophysiologist Zotterman (7) investigated cutaneous C-LTM afferents in cats using a recording electrode and biological amplifier to measure nerve activity directly in response to a stimulus applied to the nerve’s receptive field. The procedure involved dissecting the nerve out of the skin and holding it in position using a Ringer’s Solution soaked sponge, where an electrode recorded electrical activity in the nerve. Zotterman’s method was further refined for minimally invasive human applications with the development of microneurography which used etched tungsten microelectrodes and specialised amplifiers enabling the recording of single unit responses (10). Controversy surrounded the early development of the technique: firstly, it was doubted that the electrode tip would be small enough to record single-unit signals, secondly, the electrode would electrically short-out the nerve’s electrical signal and thirdly, due to the electrode manipulation, most fibres would be pressure blocked and only a few fibres would be capable of conducting a signal. These criticisms were all refuted experimentally (11).

Johansson and Vallbo (12) classified their LTM nerve recording by the *Adaptation rate* (*AR*), where nerves responding only to stimulus onset and withdrawal were characterised as fast adapting (FA) and those responding to sustained stimulation were described as slowly adapting (SA). Furthermore, nerves were also classified by *Receptive field* (RF), which describes a skin region that, when stimulated with the appropriate stimulus, evokes depolarisation in the connected afferent nerve fibre (13). RF size is dependent on the number of mechanoreceptors innervating a nerve fibre (14), and relatively small and large receptive fields are referred to as type I and II fibres, respectively (13).

Discriminatory touch, such as sensing differences in the properties of an object’s surface texture, is coded specifically by LTMs, which respond to <5mN of pressure (15). The glabrous (non-hairy) skin, such as the palm of the hand, is innervated by four types of LTM; 43% are Meissner Corpuscle (MC), 13% are Pacinian Corpuscle (PC), 25% are Merkel cell and 19% are Ruffini Corpuscle (RC) (16, 17). Due to the absence of MCs, hairy-skin is anatomically distinct from glabrous skin (10, 18).

Microneurography studies found FAI nerve fibres terminate with Meissner corpuscle endorgans, which unevenly distributed in the hand (mainly the fingertips), with a RF diameter of 2-8mm and respond to indentation forces of 0.07–5.6 mN (19–21). FAI nerves are sensitive to low frequency (2-40Hz) skin vibration, which detect slip when gripping an object (22, 23). In contrast, FAII fibres terminate with Pacinian corpuscles, which have a RF several centimeters in diameter and detect high frequency vibration from 40 to 500Hz, typically generated by object textures smaller than 1μm (22, 24, 25). Interestingly, sensitivity to higher frequency vibration (~250Hz) diminishes from age 10 to 65 years, due to morphological changes and altered spatial distribution (26). However, this effect has not been observed in hairy skin (27). FAII but not FAI firing is modulated by spatial and temporal summation (28, 29). Spatial summation describes a nerve that fires in response to simultaneous mechanoreceptor stimulation (30) and temporal summation describes a decrease in activation threshold as activation rate increases (31). SAI and SAII fibres innervate Merkel cell endings that detect object edges and Ruffini corpuscles that detect skin stretch, respectively (15, 32). However, since neither respond specifically to vibration propagated perpendicular to the skin surface, they are outside the scope of the present study.

In contrast to the discriminatory touch system, Zotterman (7) reported responses from a class of slowly conducting, unmyelinated, C-fibre LTM in cats – C-LTMs. It was initially thought that any human equivalent had disappeared through evolution, until microneurography found C-LTMs in human hairy skin sites (33, 34). Interestingly, C-LTMs have only been observed in hairy skin (33, 35). These nerves are classified as a component of the slow touch system due to a slow conduction velocity of ~1 m/sec and a 1μm diameter unmyelinated axon (1, 36, 37). C-LTMs are free nerve endings, unlike the encapsulated end organs of MTM, as described earlier, and are exquisitely sensitive to gentle stroking touch stimuli delivered at skin temperature, with an indentation force between 0.3–2.5 mN. Furthermore, C-LTM response to stimulus velocity is described by an inverted-U function, with an optimal velocity of 3cm/s and a refractory period of 30 seconds (36, 38–40).

C-LTMs are hypothesised to code for rewarding social touch because of their response properties and C-LTM activation is positively correlated with subjective pleasantness, *the social touch hypothesis* (40, 41). Pawling et al. (42) utilised electrocardiogram (ECG) and electromyography (EMG) of the zygomaticus major to investigate implicit affective response to C-LTM-optimal stimulation applied with a soft brush. Activation of the zygomaticus major was significantly greater in response to C-LTM-optimal stimulus on the arm compared to on the palm. In contradiction, no difference was found between the hairy and glabrous skin for heart rate deceleration, a proxy for measuring C-LTM activation. Pleasantness ratings of C-LTM-optimal stimuli are also rated as more pleasant on the palm compared to the arm (43).

A reported benefit of C-LTM touch, for infants, is reduced stress hormone production (44), whereas, touch deprivation increases socio-cognitive deficits (45) and increases autism rates four-fold (46). Children receiving a shoulder massage showed improved scores on abstract reasoning tests (47), whereas adults improved on accuracy, speed and alertness on a maths test (48). Furthermore, C-LTM touch is suggested to increase positive mood (49) and decrease hospital patients’ reported stress (50). McGlone et al. (51) argue that in order to flourish throughout adulthood, physical contact with others is essential and with diminished opportunities of social contact, this need is increased in the elderly. Despite the benefits, affective touch research has not received deserved attention due to the over sexualisation of normal social touch, for example, concerning foster carers (52) and schools (47).

Most current research into touch involves the development of technological innovations such as precision actuators and ultrasound haptic devices, in an attempt to artificially generate skin sensations using vibration (6, 16, 53, 54). Similar to physical C-LTM-optimal touch, pleasantness ratings of low intensity vibrotactile stimulation via actuators (200Hz) placed on the lateral forearm conformed to an inverted U-curve (40, 55). Furthermore, a similar actuator applied to the leg was capable of modulating muscle pain (56), suggesting that C-LTMs may be stimulated artificially with vibrotactile stimuli, although this has not yet been tested using microneurography (57).

In relation to the discriminatory touch system, Katz (57) demonstrated that participants were able to distinguish object textures of a grooved plate with the same accuracy when using their fingertip directly and also with passive touch when exploring the plate via a handheld tool. The researcher suggests that the mechanoreceptors react to passive stimulation as if they were directly interacting with the stimulus, Johnson (22) suggests this is due to the high sensitivity of PC afferents to small vibrations.

Many devices have been developed that deliver vibration to the skin via actuators, which have been able to create haptic feedback of a key press sensation on a virtual keyboard (51, 54) and create a pulling and pushing force sensation against the skin (58). Subjective intensity ratings of 200Hz vibrotactile stimuli delivered by a Perspex probe were found to be equal between hairy skin (dorsal forearm) and glabrous skin (fingertip) when skin deformation of the forearm was 200um more than the fingertip (18). The grounded nature of these devices creates a limitation in their use as they must be in direct contact with the skin and the attached wiring is an inconvenience to free movement.

Overcoming this limitation, a Japanese research team developed a novel ungrounded stimulator, known as an ultrasonic mid-air haptic device (18). This comprises a lattice of ultrasound transducers, also referred as a phased array, where each transducer can be individually controlled via computer software. Such control enables the converging of emitted sound waves at one (or more) defined point(s) in space, also referred to as focal point(s), where acoustic pressure is significantly higher than its surrounding (59). Due to the non-linear phenomenon arising from second order perturbation, so-called Acoustic Radiation Pressure (ARP) is produced. Effectively, ARP is a non-zero time-average force being exerted on a surface and is roughly proportional to the square of the acoustic pressure. With commercially available devices, such as the one produced by Ultraleap, ARP can reach up to 1gf. When applied to the skin, ARP can deform the skin up to a few micrometers (60), and therefore has the potential to activate cutaneous mechanoreceptors and induce tactile sensation.

Ultrasound mid-air haptic devices commonly use 40kHz ultrasound transducers. The effective size (or width) of the focal point is inversely proportional to the transducer frequency, and therefore using 70kHz ultrasound transducers to produce smaller focal points has been suggested (61). In addition to be inaudible, both frequencies are way beyond the sensitivity range of mechanoreceptors and therefore the force applied by the focal point on the skin can be assumed sustained. Since MC and PC receptors do not depolarize to sustained force, it has been proposed to incorporate vibratory behaviour within the sensitivity range of MC and PC receptors through two modulation techniques: Amplitude Modulation and Spatiotemporal Modulation (62).

Amplitude Modulation (AM) describes a method of temporarily changing the amplitude of the focal point acoustic pressure. Specifically sinusoidal waveform, at frequency within the range of MC and PC receptors, are used to induce vibrotactile effect. Commonly a modulation of 200Hz is used as Brisben et al. (63) found utilising a vibrating cylinder and detection threshold of vibration in the hand varied as a function of vibration frequency and was described by a U-shaped function, where 200Hz was the lowest detectable stimulus threshold. Raza et al. (64) found a similar function with mid-air haptics AM stimuli and suggest that ARP should be tuned with AM frequency to ensure all AM stimuli to be perceived equally strong. Fan et al. (65), also showed that using modulation waveform with different Root-Mean Square, different measure of ARP could be observed on a precision scale. This is expected, as modulation waveform are applied on the acoustic pressure, and ARP is proportional to a time-average of the acoustic pressure square. So while the time averaged values are different, the peak ARP with all waveform remain the same. Besides perceived strength, Obrist et al. (24) found that the overall sensation produced by a mid-air haptic device vary with the modulation frequency. As such, an AM stimulus is described as “puffs of air” at 16Hz (MC range) and as a “breeze” at 200Hz modulation (PC range).

Spatiotemporal Modulation (STM) describes a method of temporarily changing the position of the focal point, while amplitude is kept constant. Specifically, the focal point repeats closed curve at a rate fast enough for the user to perceive the closed curve as a continuous shape, a phenomenon similar to the persistence of vision. While the ARP of STM pattern is kept constant through the pattern, Frier et al. shows that the amount of displacement produced could vary with the speed of the focal point along said pattern (62). Authors, also show that this variation of displacement were positively correlated with perceived strength. Takahashi et al. (66) were the first to investigate the detection threshold and subjective perception of both AM and STM stimuli. They utilised a custom-built haptic device emitting 40kHz ultrasound downwards onto the participants’ arms in a small-scale study. The detection threshold of STM stimuli was significantly lower than AM stimuli on both glabrous and hairy skin. Furthermore, vibrotactile STM stimulation on the hairy skin of the arm was rated as more intense than AM.

The application of mid-air haptic technology is varied and requires multi-disciplinary collaboration between cognitive neuroscience and affective computing (67). The addition of gesture tracking technology (“Leap Motion”) to mid-air haptic devices (“Stratos Inspire”), facilitates targeting specific skin regions on the hand and fingers (6). The combination of both technologies have lead researcher in Human Computer Interaction (HCI) to explore novel interfaces, which have been compiled in a recent survey by Rakkolainen et al. (67).

Most literature describes affective touch as a necessity for the development of cognitive processing and not an indulgence (45, 46). However, given the increasing taboo of social touch (47) and increased global social isolation resulting from the Covid-19 pandemic measures (68), the potential to develop an ultrasound alternative for social touch, may in the future, prove therapeutic for those deprived of physical touch. In the case of virtual reality (VR) and digital interfaces, ultrasound could provide a way of interacting with the digital virtual world as if it were physically present, facilitating computer-human interfacing (69). However, little is known about subjective intensity and pleasantness ratings of ultrasound stimulation and the bottom-up and top-down factors mediating the effects. Furthermore, microneurography has never measured mechanoreceptor activation during ultrasound stimulation.

The present study aims to (1) investigate subjective intensity and pleasantness ratings of ultrasonic skin stimulation and the influence of top-down factors, and (2) use microneurography to determine which classes of mechanoreceptors are activated by ultrasound stimulation of the skin.

It is hypothesised that (1) There will be a significant three-way interaction for intensity ratings between skin type, output power and modulation; (2) Intensity ratings will be significantly different between age groups (above and below 24 years); (3) Intensity ratings will be significantly correlated with TEAQ sub-scale scores; (4) There will be a significant three-way interaction for pleasantness ratings between skin type, output power and modulation; (5) Pleasantness ratings will be significantly different between age groups (above and below 24 years); (6) Pleasantness ratings will be significantly positively correlated with TEAQ sub-scale scores.

## Method

### Participants

Fifty-one participants recruited via opportunity sampling, had an average age of 26.33 years (SD = 9.77) and twenty-five were male. Exclusion criteria included anyone under 18 years; or those with a skin condition, neurological condition, heart conditions, pacemaker; and those who are pregnant or elderly.

### Design

The present study was a 2×2×2 within-subjects group design utilising two 3-way repeated measures ANOVA.

### Materials

#### Haptic Array

The Stratos Inspire device, consisting of 256 ultrasonic transducers emitting 40kHz sinusoidal sound waves. A custom BASH script executed a customised Stratos Inspire C++ API. This software allowed for the creation of a mid-air haptic focal point at a distance of 20cm set to two output powers (pascal scale), 100% (163dB) and 50% (158dBs) with two modulations (AM and STM). The haptic device was mounted upside down onto a stand and two low-powered lasers were attached to visualise the focal point.

#### Chair and Suction Cushion

A dentist style chair was half reclined and had two arm rests. A suction cushion was moulded to the participants left arm, minimising movement.

#### Scientific Scales

The analytical balance (Acculab ALC80.4) had an 80mm diameter plate, a glass housing and a 0.0001g resolution.

#### Demographic Questionnaire

A two-item demographic self-report recorded participants age and gender.

#### TEAQ

The Touch Experience and Attitudes Questionnaire (TEAQ) is a 57 item self-report employing a 5-point Likert scale from “Disagree strongly” to “Agree strongly”. The TEAQ comprised six sub-scales, namely; current social touch (CST), current intimate touch (CIT), childhood touch (ChT), attitude to self-care (ASc), attitude to intimate touch (AIT), and attitude to unfamiliar touch (AUT). Disagree strongly was scored as one and agree strongly was scored as five, except for items 1, 3, 9, 23, 28, 37, 39, and 53, which were reverse scored. Sub-scale mean scores were calculated for each participant.

#### Visual Analogue Scale (VAS)

Intensity and pleasantness ratings were recorded using two 100mm VAS. The intensity scale ranged from 0-100. The pleasantness VAS ranged from −10 to +10 and scores on both scales were calculated by measuring the distance of the rating from zero.

#### Microneurography

The microneurography equipment consisted of an uninsulated subdermal reference electrode and an epoxy-resin insulated 40mm tungsten electrode, with a 5μm diameter bare tip. A high-impedance preamplifier was connected to a low-noise high-gain amplifier, which connected to a computer were the nerve signal was displayed by LabChart and heard through a speaker. Von-Frey hairs (0.1-100g), a tuning fork (125Hz) and soft and hard brushes were used to categorise the nerve fibre.

### Procedure

#### Experiment 1: VAS

Utilising an opportunity sampling method, LJMU students were recruited via an online booking system (SONA) and invited to the psychology lab. Participants read a participant information sheet before completing a consent form, demographic questionnaire and the TEAQ. Finally, participants sat in the chair with their left palm up, supported on a suction cushion. The experimenter’s hand was placed under the focal point and lasers were aimed at the point of sensation. The haptic device position was adjusted so that the focus point targeted the participant’s palm (fig 1). Ear defenders were worn by participants during the delivery of eight stimuli, each lasting 10 seconds in the order shown in table 1, and the start and end of stimulus delivery was marked by a short sound played through the laptop speakers. A 20 second pause was provided between stimuli allowing time for participant VAS responses, therefore amounting to 4m30sec total run time. Following stimuli two and six the device was manually moved to the arm and palm, respectively.

**Fig 1.**
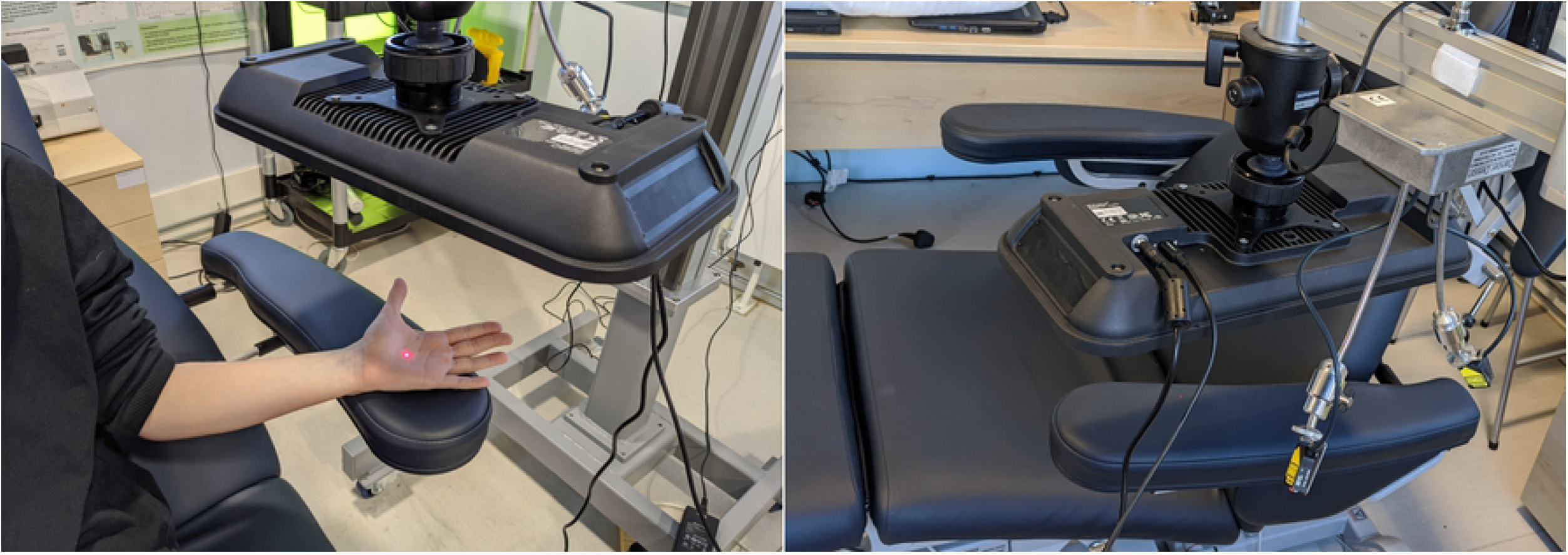
Haptic device setup. Image depicting haptic device setup with laser dot indicating the ultrasound focal point on the participants palm (left); and laser housing mounting (right).

**Table 1.** Stimulus protocol comprising eight stimuli varying in skin location, device output power and modulation.

| Stimulus | Skin<sup>a</sup> | Output Power<sup>b</sup> | Modulation<sup>c</sup> | Pattern | Frequency | Duration |
| --- | --- | --- | --- | --- | --- | --- |
| 1 | Palm | 100% | STM | 2cm Circle | 10cm/s | 10 seconds |
| 2 | Palm | 50% | STM | 2cm Circle | 10cm/s | 10 seconds |
| 3 | Arm | 100% | STM | 2cm Circle | 10cm/s | 10 seconds |
| 4 | Arm | 50% | STM | 2cm Circle | 10cm/s | 10 seconds |
| 5 | Arm | 100% | AM | Point | 100Hz | 10 seconds |
| 6 | Arm | 50% | AM | Point | 100Hz | 10 seconds |
| 7 | Palm | 100% | AM | Point | 100Hz | 10 seconds |
| 8 | Palm | Low | AM | Point | 100Hz | 10 seconds |

<sup>a</sup> Arm = hairy skin of the dorsal forearm and palm = centre of left palm
<sup>b</sup> 100% output power = 163dB and 50% output power = 158dB.
<sup>c</sup> STM = spatiotemporal modulation and AM= Amplitude modulation.

#### Experiment 2: Microneurography

Nerve type classification was determined by inserting the electrode percutaneously, 1-3cm proximal to the cubital fold, into the lateral antebrachial nerve. Sensations down the arm produced by weak electrical pulses (0.1-3mA) indicated the electrode was interfascicular. Nerve fibre tuning characteristics and receptive field size were identified using Von-Frey hairs (<5mN), tuning fork vibration and gentle stroking touch. Nerve activation was record in response to all ultrasound outputs and modulations.

#### Experiment 3: Acoustic radiation pressure

The haptic device was mounted facing downwards above the scientific balance, with the glass housing open. The focal point was aimed (as above) at the balance plate, 40cm below and measurements were recorded.

### Ethical considerations

The present study was conducted to British Psychological Society ethical standards and received LJMU’s ethics committee approval. Although the Stratos Inspire device is considered safe via CE marking, ear defenders were worn by all persons present to protect the ears from prolonged exposure to ultrasound. Consistent with the contraindications noted in the Startos Inspire manual, participant exclusion criteria consisted of those with a; skin conditions, neurological conditions, pacemaker, implantable cardioverter defibrillator and heart conditions; or those who were pregnant or elderly (Ultraleap, 2020). To protect the participant from experiencing physical or psychological discomfort resulting from participation, a participant information sheet explained the purpose of the study, stated that questions may be skipped on the questionnaire and highlighted that participants may withdraw at any time, for any reason. Identifiable information was not collected to maintain confidentiality; thus, withdrawal of data was not possible due to data being anonymised. Informed consent was gained in writing via a consent form. A debrief sheet provided a detailed explanation of the aims and hypotheses of the study. Furthermore, to protect the participant from any harm following the study, contact details of the researcher and study supervisor was provided. Reducing the risk of physical discomfort or harm resulting from multiple microneurography sessions, participants were only permitted to take part in the study once.

### Analysis

Outliers were excluded from analysis based on z-scores more than +/- 2, thus, 11 extreme values were identified based on pleasantness scores and 16 for intensity scores. A 2×2×2 repeated measures ANOVA was conducted.

## Results

Table 2 indicates that stimulus seven was rated the most intense and pleasant and had the largest standard error, whereas, stimulus six was rated the least intense and pleasant, and had the smallest standard error. Both intensity and pleasantness ratings of the arm (stimuli 3-6) were generally reported with higher scores and smaller standard errors compared to stimuli on the arm.

**Table 2.** Means and standard errors of intensity and pleasantness ratings for all eight stimuli varying on skin location, device output power and modulation.

| Stimulus | Intensity rating |  | Pleasantness rating |  |
| --- | --- | --- | --- | --- |
|  | M | SE | M | SE |
| 1<sup>a</sup> | 9.97 | 1.64 | 1.35 | 0.27 |
| 2<sup>b</sup> | 13.18 | 2.10 | 1.21 | 0.27 |
| 3<sup>c</sup> | 8.37 | 1.45 | 0.76 | 0.19 |
| 4<sup>d</sup> | 11.60 | 2.06 | 0.56 | 0.19 |
| 5<sup>e</sup> | 7.63 | 1.79 | 0.35 | 0.24 |
| 6<sup>f</sup> | 2.96 | 0.89 | 0.18 | 0.18 |
| 7<sup>g</sup> | 56.03 | 3.70 | 3.22 | 0.44 |
| 8<sup>h</sup> | 30.97 | 4.71 | 1.56 | 0.31 |

<sup>a</sup> high output STM on palm, <sup>b</sup> low STM on palm, <sup>c</sup> high STM on arm, <sup>d</sup> low STM on arm, <sup>e</sup> high AM on arm, <sup>f</sup> low AM on arm, <sup>g</sup> high AM on palm, <sup>h</sup> low AM on palm. Both perceived intensity and pleasantness scores were reported as greatest and lowest for stimuli seven and six, respectively.

A Kolmogorov-Smirnov (KS) test assessed normality of data. Intensity ratings were not normally distributed, Ds(35) < .36, *p* < .05, except for *high output AM on the palm*, D(35) = .07, *p* > .05. Supporting hypothesis one, a significant three-way interaction was found between skin, output and modulation, *F*(1,36) = 13.70, *p* < .05, ηp^2^ = .29. A 2×2×2 repeated-measures ANOVA analysed intensity and pleasantness data (fig 2 & 3). A significant one-way interactions was found for intensity ratings between; the palm (*M* = 27.54, *SE* = 2.34) and arm (*M* = 7.64, *SE* = 1.04), *F*(1, 24) = 81.4, *p* < .05, ηp^2^ = .71; low (*M* = 14.68, *SE* = 1.69) and high output (*M* = 20.5, *SE* = 1.39), *F*(1, 34) = 25.04, *p* < .05, ηp^2^ = .42; however, no significant difference was found between AM (*M* = 24.39, *SE* = 2.13) and STM (*M* = 10.78, *SE* = 1.22), *F*(1,34) = 47.78, *p* > .05, ηp^2^ = .58. A Wilcoxon signed-ranks test indicated that intensity ratings were significantly different between high and low output for AM stimuli (Zs < 4.70, *ps* < .05), but not for STM stimuli (Zs < 1.77, *ps* > .05). Wilcoxon signed-ranks tests showed that intensity ratings of AM were significantly higher on the palm than the arm, Zs < 5.7, *ps* < .05, but not for STM Zs < .79, *ps* < .05.

**FIG 2.**
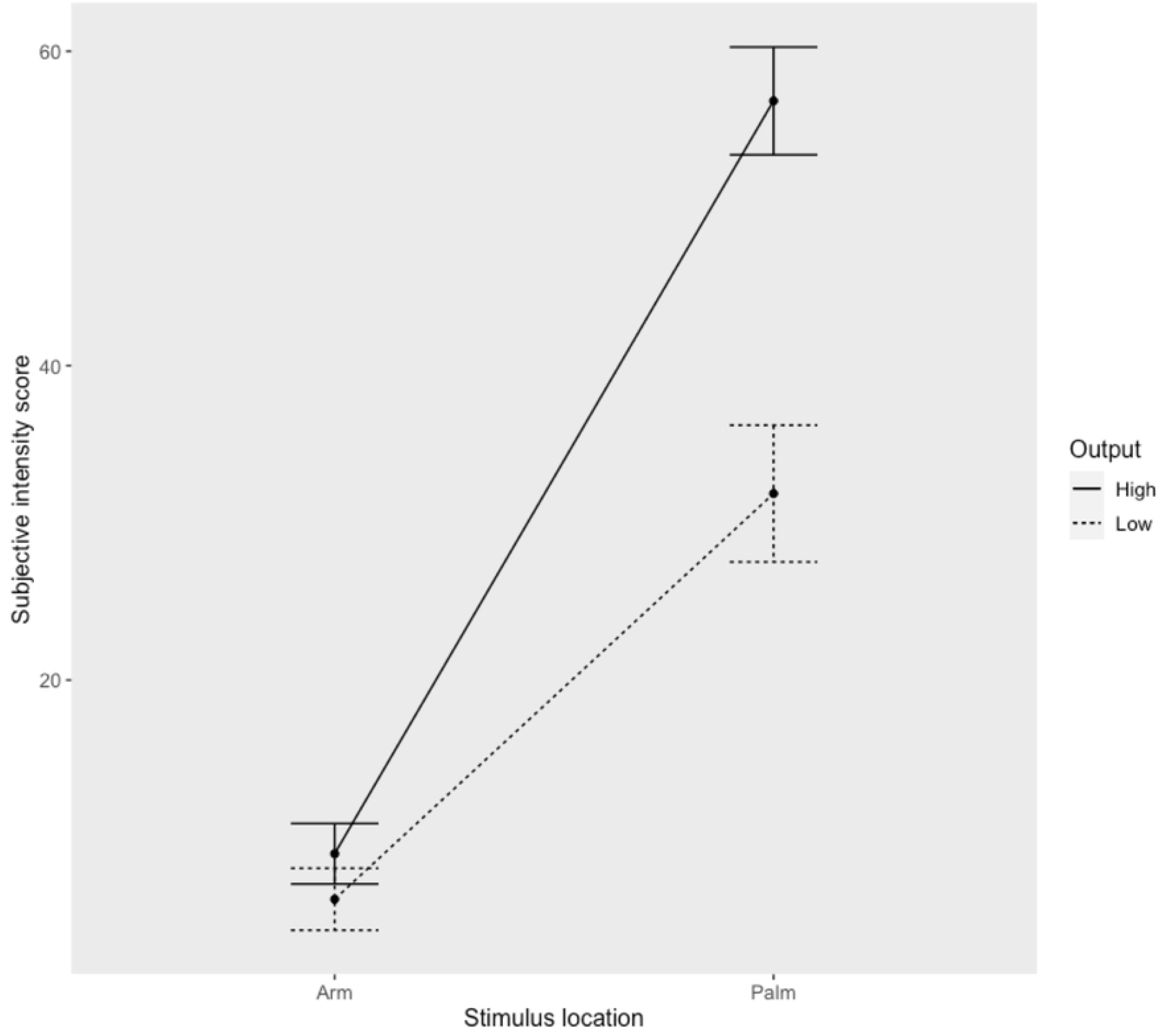
Intensity ratings for AM ultrasound stimuli as a function of high (100%) and low (50%) device output on the palm and hairy skin of the forearm. For both output levels, subjective intensity was reported as greater on the glabrous skin of the palm compared to the hairy skin of the forearm. Error bars indicate standard deviation.

**FIG 3.**
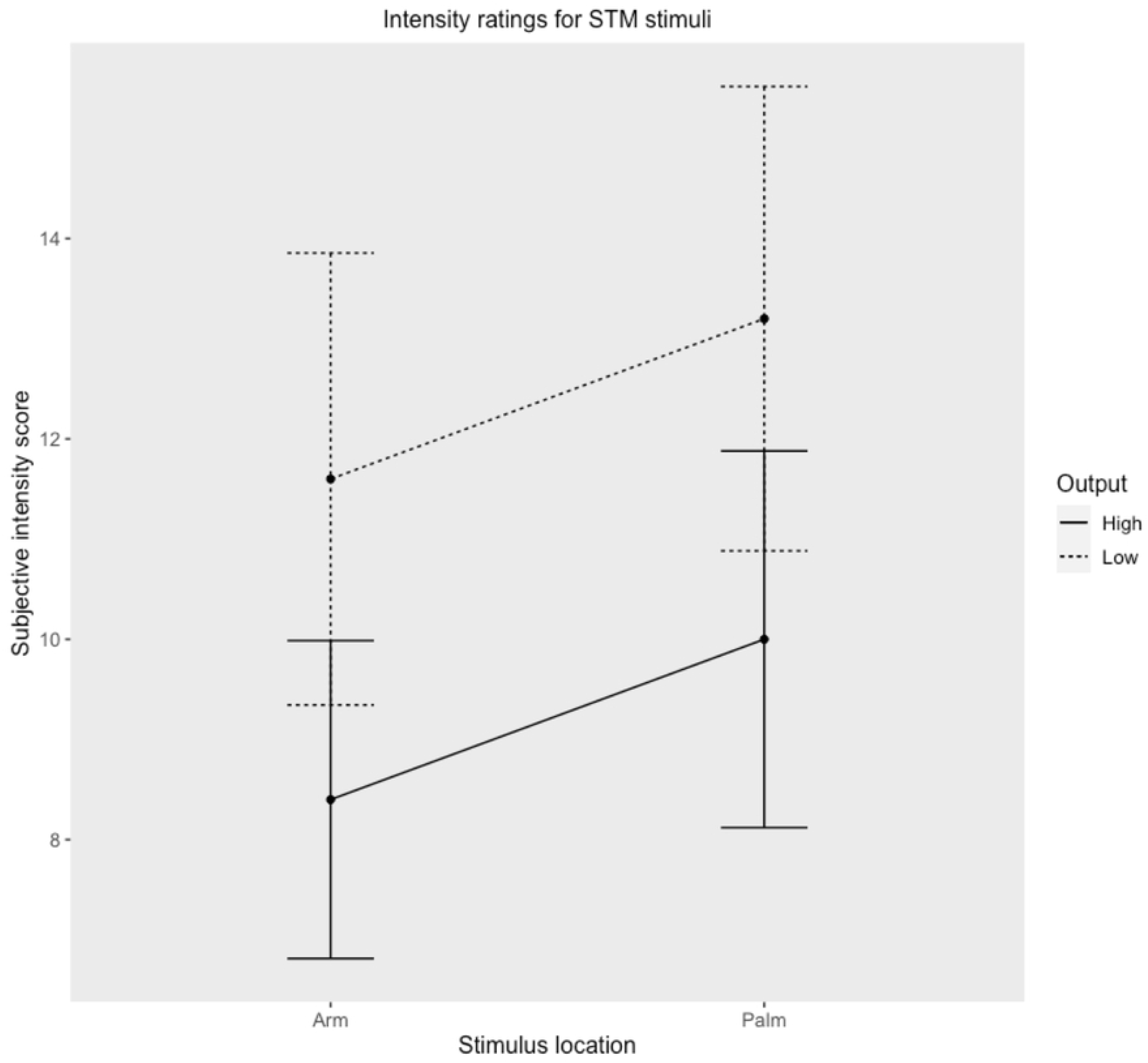
Intensity ratings for spatiotemporal modulation ultrasound stimuli as a function of high (100%) and low (50%) device output on the palm and hairy skin of the forearm. For both output levels, subjective intensity ratings were greater for the glabrous skin of the palm compared to the hairy skin of the forearm. Error bars indicate standard deviation.

A K-S test found intensity rating data for the under 25 years group to be normally distributed for stimuli 1-4, 7-8, Ds(29) < .15, *ps* > .05, but not for stimuli 5-6, Ds(29) < .30, *ps* < .05. For the over 25 years group, data was normally distributed for stimulus 7, D(15) = .20, *p* > .05, but not for stimuli 1-6 and 8, Ds(7 < .39, *ps* < .05. Supporting hypothesis two, a Mann-Whitney U test found intensity ratings for stimuli 1-3 and 8 to be significantly higher in the younger group compared to older group, Us < 125.00, *ps* < .05 (fig 4).

**FIG 4.**
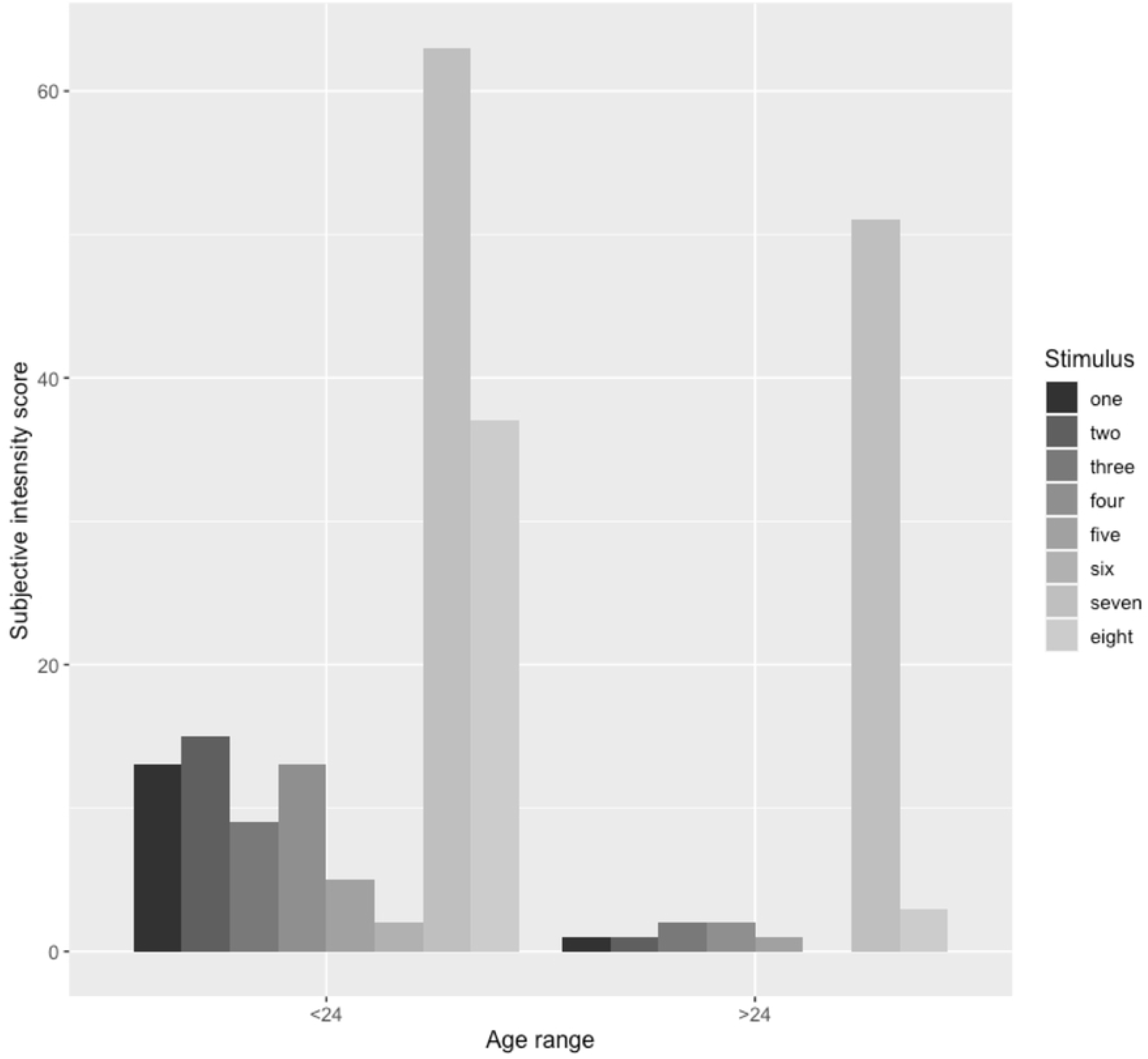
Intensity ratings between age groups (<24 years & >24years) for all stimuli, described by location (arm/palm), device output (high/low) and modulation type (spatiotemporal modulation/amplitude modulation). For the younger age group, subjective intensities were reported as greatest for stimuli seven and eight, whereas for the older age group, subjective intensity was reported as greatest for stimulus seven.

Rejecting hypothesis three, a Pearson’s correlation found that intensity ratings were not significantly correlated with the six TEAQ subscales, *rs*(51) < .24, *p*s > .05.

For pleasantness ratings, a KS test showed data were not normally distributed, Ds < .28, *p* < .05. Supporting hypothesis four, a 2×2×2 repeated measures ANOVA found a significant threeway interaction between skin, output and modulation, *F*(1, 39) = 6.98, *p* < .05, ηp^2^ = .15 (figure 5 & 6). Significant one-way interactions for pleasantness ratings were found between; the palm (*M* = 1.83, *SE* = .26) and arm (*M* = .46, *SE* = .14), *F*(1, 39) = 43.57, ηp^2^ = .53; Low output (*M* = .87, *SE* = .17) and high output (*M* = 1.42, *SE* = 1.75), *F*(1, 39) = 10.29, *p* < .05, ηp^2^ = .21; AM (*M* = 1.32, *SE* = 1.85) and STM (*M* = .97, *SE* = .17), *F*(1, 39) = 3.23, *p* < .05, ηp^2^ = .07; A Wilcoxon signed-ranks test indicated that pleasantness ratings were not significant difference between high and low output for AM (Z = .15, *ps* > .05), except for between stimulus seven and eight (Zs = −3.03, *ps* < .05) (fig 5). High and low output were not significantly different for STM stimuli (Z < .88, *ps* > .05) (fig 6).

**FIG 5.**
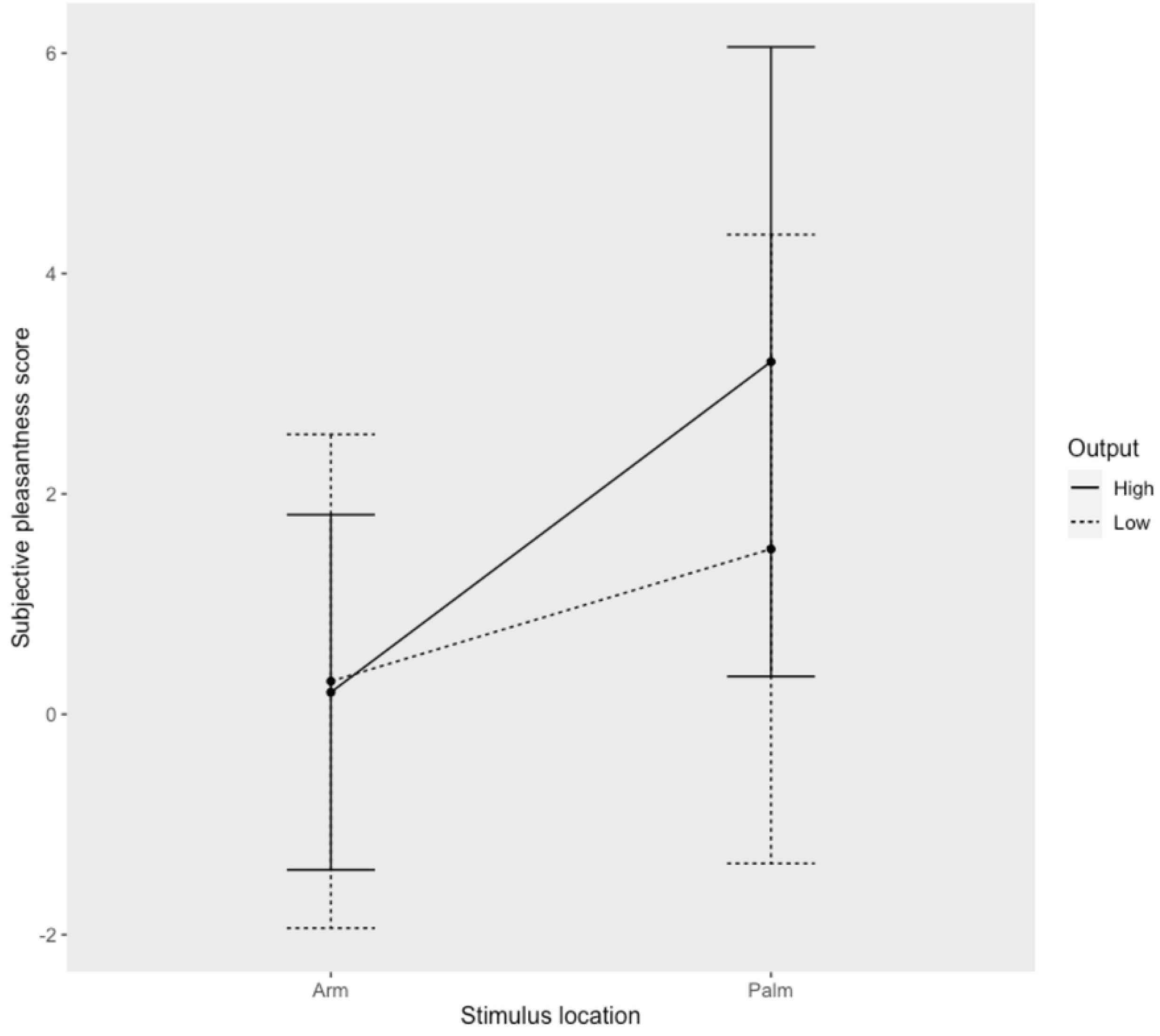
Pleasantness ratings for amplitude modulated ultrasound stimuli as a function of high (100%) and low (50%) device output on the palm and hairy skin of the forearm. Error bars indicate standard deviation. For both output levels, subject pleasantness ratings were greatest for the glabrous skin of the palm compared to the hairy skin of the forearm. Error bars indicated standard deviation.

**FIG 6.**
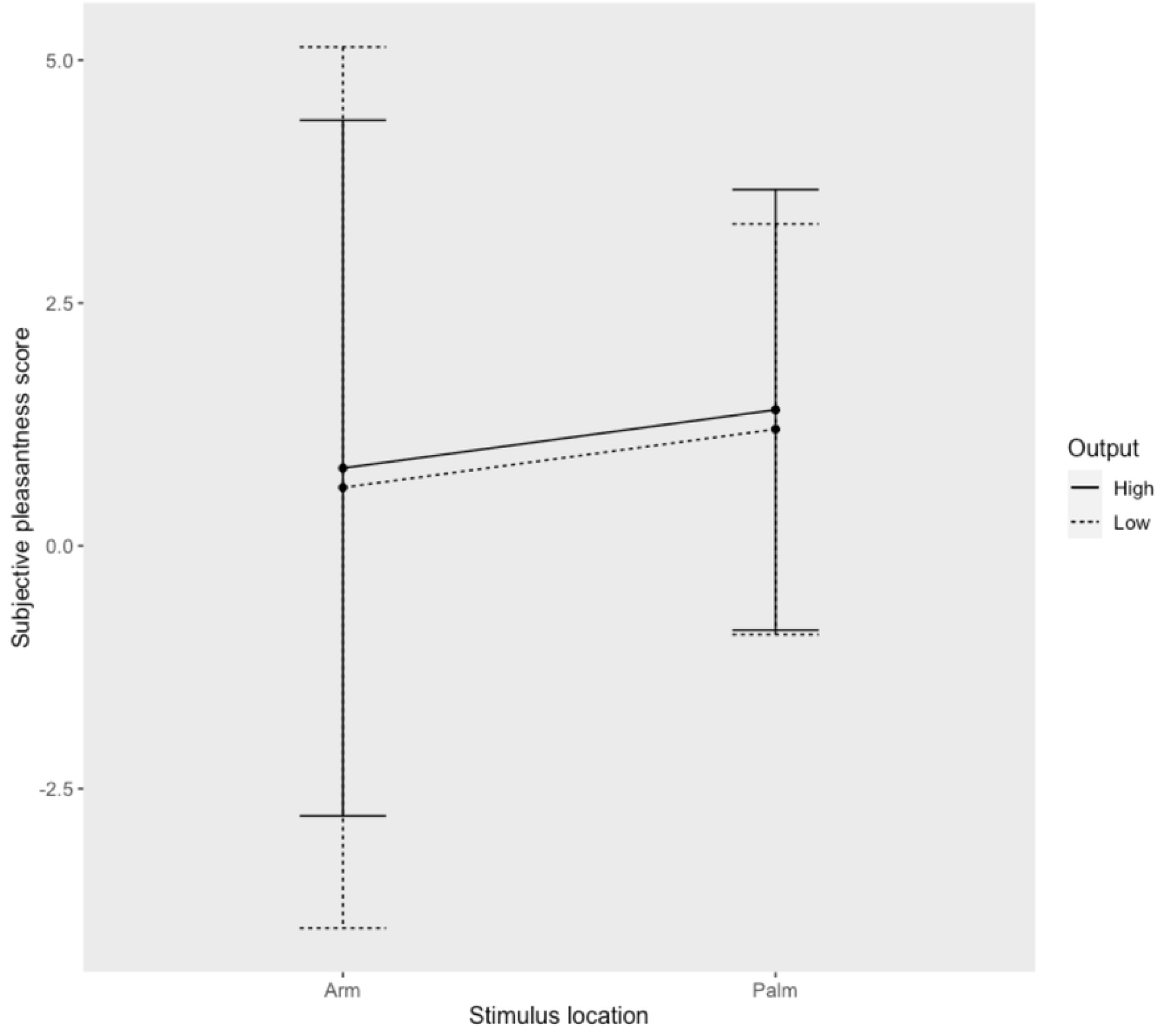
Pleasantness ratings for spatiotemporal modulated ultrasound stimuli as a function of high (100%) and low (50%) device output on the palm and hairy skin of the forearm. Error bars indicate standard deviation. For both output levels, subject pleasantness ratings were greatest for the glabrous skin of the palm compared to the hairy skin of the forearm. Error bars indicate standard deviation.

For the younger group, a K-S test found data to be normally distributed for stimuli 1-2, Ds(28) < .14, *ps* > .05, but not for stimuli 3-8, Ds(28) < .33, *ps* < .05 (fig 7). For the older group, data were normally distributed for stimulus 1-6 and 8, D(16) < .37, *p* > .05, but not for stimuli 7 (D(16) = .20, *ps* < .05). Partially supporting hypothesis five, a Mann-Whitney U test found pleasantness ratings of stimuli 1-3 to be significantly different between younger and older participants (Us < 141.50, *ps* < .05) but not for stimuli 4-8 (Us < 217.00, *p* > .05).

**FIG 7.**
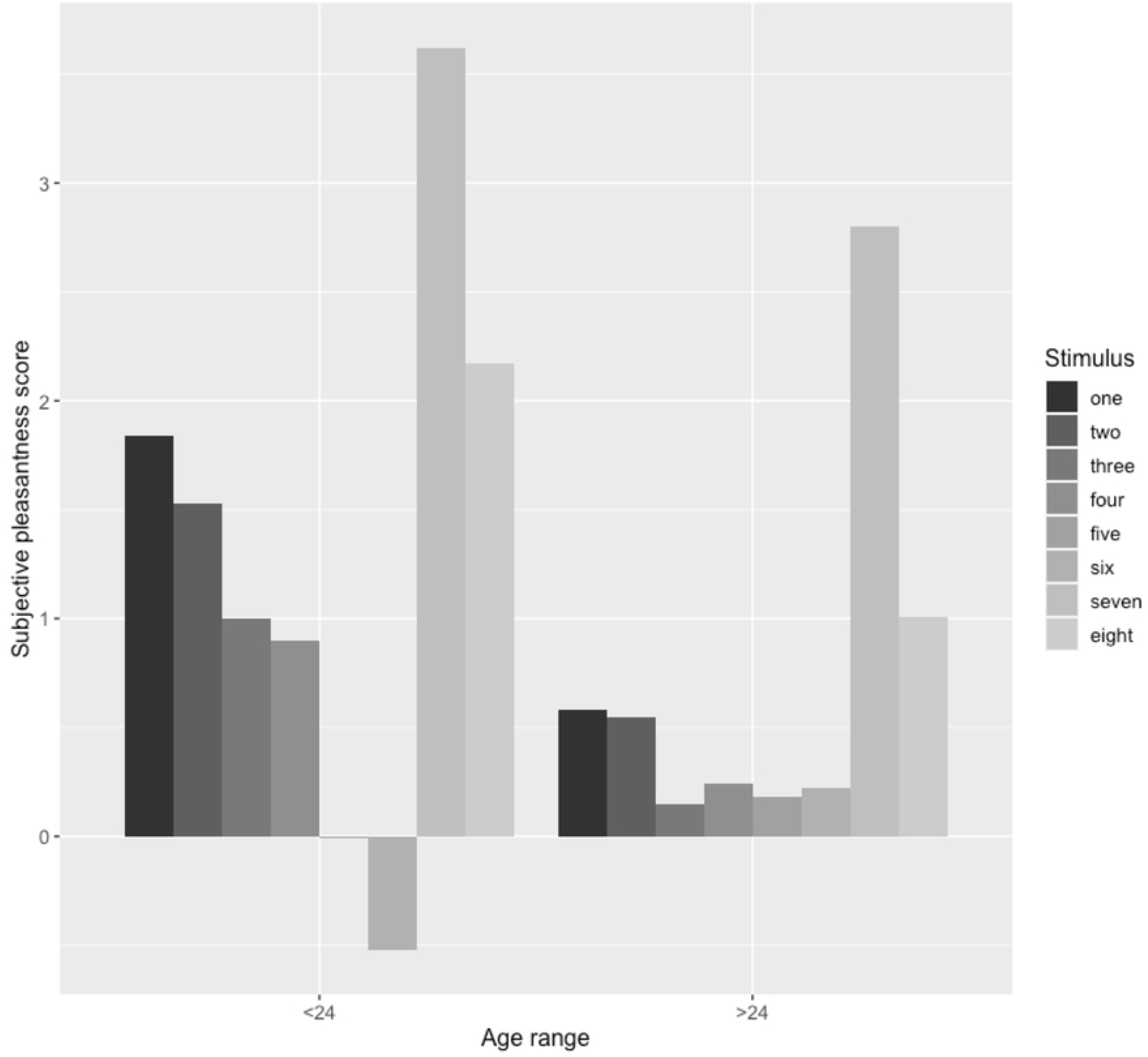
Pleasantness ratings between age groups (<24 years & >24years) for all stimuli, described by location (arm/palm), device output (high/low) and modulation type (spatiotemporal modulation/amplitude modulation). For both age groups, stimulus seven was rated as most pleasant. Generally, pleasant ratings were greater for the younger age group; however, stimulus six was the only stimulus to be rated as unpleasant, but only by the younger age group.

Partially supporting hypothesis six, pleasantness ratings were not significantly correlated with the six TEAQ subscales, *rs*(51) < .28, *p*s > .05, except for a positive correlation between stimulus four and *attitude to intimate touch*, *rs*(51) < .29, *p*s < .05.

The ARP experiment data (Table 3) shows the downwards force generated by each modulation and device output power. The data indicates AM at 100% and 50% output power produced a much lower ARP, (1.96mN and 0.39) compared to STM (7.06mN and 6.57mN).

**Table 3.**
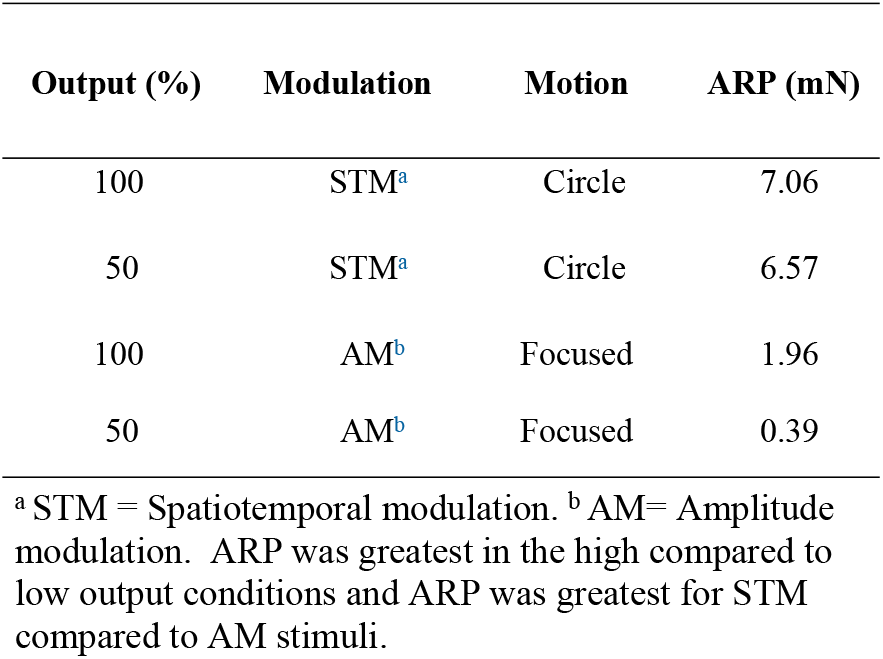
The weight (mn) of air displacement at a 20cm focal point for various output levels (ARP), modulations and motions.

| Output (%) | Modulation | Motion | ARP (mN) |
| --- | --- | --- | --- |
| 100 | STM<sup>a</sup> | Circle | 7.06 |
| 50 | STM<sup>a</sup> | Circle | 6.57 |
| 100 | AM<sup>b</sup> | Focused | 1.96 |
| 50 | AM<sup>b</sup> | Focused | 0.39 |

<sup>a</sup> STM = Spatiotemporal modulation. <sup>b</sup> AM= Amplitude modulation. ARP was greatest in the high compared to low output conditions and ARP was greatest for STM compared to AM stimuli.

Microneurography testing found that two median nerve units responded to a 1g Von Frey hold and 125Hz vibration and were classified as low-threshold, SAI and SAII nerves, which failed to respond to all ultrasound stimuli. The units were both still present following ultrasound stimuli. Therefore, contradicting the expectation of FAII and C-LTM mechanoreceptor activation in response to all ultrasound stimuli.

## Discussion

The present study aimed to investigate subjective intensity and pleasantness ratings of ultrasonic stimulation and the influence of top-down factors. Microneurography was used to determine which mechanoreceptors are activated by ultrasound stimulation of the skin. It was hypothesised that; there would be a significant three-way interaction for intensity and pleasantness ratings between skin types, output and modulation; intensity and pleasantness ratings would be significantly different between age groups (above and below 24 years); finally, intensity and pleasantness ratings would be significantly positively correlated with TEAQ subscale scores. The major findings were that STM stimuli produced more downwards force compared to AM; only intensity of AM stimulus was different between skin types; intensity and pleasantness ratings were significantly different between output power, modulations and age groups; only the TEAQ correlated with the pleasantness rating of stimulus four (C-LTM-optimal stimulus); and that microneurography found SAI and SAII units failed to respond to all stimuli; Intensity ratings were greatest for low output AM on the palm, which may be explained by AM generating forces within LTM range, 0.07–5.6mN (21), whereas STM forces far exceeded this. Despite this, STM stimuli did still produce a skin sensation, supported by positive ratings, which for the arm were similar to AM intensity ratings. This may be explained by a decreasing pressure gradient extending outwards from the focal point (60). It seems plausible that the force generated on the outer edges of the focal point, may be within LTM force range. While the peak ARP is the same between modulations, the time-average ARP is different as observed on the precision scale. One could say that this difference would explain the lower detection threshold of STM compared to AM (66, 70). However, with STM the focal point moves across the skin surface, and therefore does not stay directly above the mechanoreceptor receptive field. Therefore, if we consider solely the time-average of ARP applied to the mechanoreceptor receptive field, in both AM and STM, the difference would be far less than one would initially expect considering the ARP applied to the whole skin surface. Furthermore, equal force does not imply equal displacement on the skin as shown in previous study (62).

Intensity ratings for STM stimuli did not differ between the arm and palm, regardless of output power, suggesting the activation of a common mechanoreceptor between skin types, likely the PC, based on tuning characteristics (18, 28). However, high and low output AM stimuli were rated approximately 10 times and 7 times more intensely on the palm compared to the arm, respectively. Greater palm intensity ratings are consistent with the literature (18) and may suggest that a class of mechanoreceptors are being activated in the palm, which are absent in the arm. Based on tuning properties, the most likely mechanoreceptor is the MC (18, 28).

The present study demonstrates for the first time, that age modulates intensity ratings of ultrasound stimuli, where the younger group rated STM (stimuli 1-3) and AM (stimulus seven) higher on both the glabrous and hairy skin. Age effects are associated with the decline in PC mechanoreceptor sensitivity to vibrotactile stimuli (26), although no age effects have also been reported (27). Confusingly, stimuli 1-3 were STM modulated and as discussed above, were outside PC force range, but the associated age effects were still demonstrated.

The order of stimuli, based on highest to lowest pleasantness rating, is similar to the order of intensity ratings, this is also true based on output level and between age groups. Since pleasantness of ultrasound is not described in the literature, and the present study did not control for subjective intensity, it is difficult to speculate on the reason for differences between pleasantness ratings. This may suggest that higher intensity stimuli were more pleasant simply because the stimulus was more salient.

Pleasantness ratings contradicted the social touch hypothesis (40, 41) as stimulus four (low output STM) on the arm was rated as the least pleasant and stimulus 7 (high output AM) on the palm was the most pleasant. However, this is consistent with vibrotactile literature, where the glabrous skin was rated more pleasant than the C-LTM-innervated hairy skin (42, 43).

Pleasantness ratings of low output STM on the arm (stimulus four that was designed to be approximately C-LTM-optimal) were positively correlated with *attitude to intimate touch scores* (*TEAQ*). The literature supports this finding as social touch predominantly involves C-LTM-optimal touch (40, 41). Despite this correlation, it seems unlikely that C-LTM activation occurred, for four reasons: Firstly, stimulus four is fundamentally different from gentle stroking touch. Secondly, stimulus four generated an ARP (6.57mN) exceeding the C-LTM-force threshold (0.3-2.5mN) (36). Thirdly, C-LTMs are tuned to skin temperature (38), however, skin deformation was generated with ultrasound compressing air at room temperature. Fourthly, although not empirically tested, it seems plausible that wind-chill effect, caused by air moving over the skin (71), caused localised cooling, attenuating C-LTM activation.

Microneurography recorded nerve fibre responses to ultrasound stimulation from two SAI and SAII units, which failed to respond to all stimuli. Although this finding was expected, as slowly-adapting fibres respond to skin stretch only (15, 32), it is in itself a novel finding, as microneurography has never before shown that slowly-adapting units do not respond to specific ultrasound stimuli. It was expected that FAI and FAII fibres would respond to ultrasound due to their tuning properties (1, 22, 24, 25) and future studies will aim to record FA fibres.

The downwards force (ARP) of each stimulus was measured using a precision balance (Table 3). Higher output power generated a greater ARP upon the skin for all stimuli. Overall, STM produced a significantly greater ARP compared to AM, which is consistent with STM displacing the skin surface more than AM (60). The result difference between modulations was expected due to the difference in focal point amplitude over time and the fact that precision scales measure the time average of ARP and not the peak ARP. Indeed, while amplitude stays constant with STM (i.e. at maximum level), with AM the focal point amplitude fluctuates and the associated ARP has a time average lower than with STM.

The present study had a number of limitations affecting internal and external validity. Firstly, to visualise the position of the focal point in mid-air, two lasers were calibrated by aiming them at the point on the researcher where a stimulus was felt. This method was intended to maintain a 20cm distance between the skin surface and the device, producing a consistent stimulus between skin locations and between participants. However, relying on a subjective measure of the focal point position may not be an accurate protocol and could have led to stimuli being administered at different distances above the skin, therefore affecting the degree of skin deformation and in turn sensation produced. The present method would benefit from a mounting rig, where the lasers are fixed in position to standardise the point of intersection and therefore the focal point position. This would facilitate delivering stimuli at a consistent distance from the skin. This study attempted to compare intensity ratings between different modulations, skin types and output levels of ultrasound.

However, experiment three highlighted the difference between peak ARP, which is the same between modulations, and time-average ARP, which can be either, different when considering the force applied to the skin, or analogous when considering the force applied to a single mechanoreceptor receptive field. Due to this ambiguity regarding time-averaged ARP, this study controlled for the peak force generated. Nonetheless, the possibility remains that time-average ARP on the skin is the quantity actually perceived by C-LTM, and that STM stimuli did not produce C-LTM-optimal forces range (36). Due to this comparison between differences in subjective intensity and pleasantness ratings should be done cautiously. Similar carefulness should be applied when considering the external validity and generalisability of the correlation between *attitudes to intimate touch* (TEAQ) and pleasantness ratings of stimulus four. The issue of ARP difference may be addressed by including more stimuli with controlled peak ARP as well as controlled time-average ARP. With the same line of thoughts, one may attempt to control for skin displacement on the skin.

Finally, an opportunistic sampling method did not provide a representative sample of the population, due to 61% of the sample being aged between 20-21 years. Therefore, the generalisability of the age-related data is reduced. Furthermore, since data was not collected from participants under 21 years of age, intensity and pleasantness ratings of ultrasound for children and adolescents remain untested. This study would benefit from a stratified sample, recruiting participants from various age ranges.

The implications of the reported findings involve the haptics industry. Reporting that STM was rated with equal intensity on the palm and arm, has implications for VR, where producing sensations with consistent intensity between skin types is desirable. For example, virtual posters may take advantage of this, allowing the user to feel the texture of a dress, with a consistent sensory experience, whether the virtual item is held in the hand or draped over the arm (72). In terms of haptic device users, the suitability of ultrasound may be more appropriate for those under 24 years of age, since stimulus intensity decreases with age. However, age effects did not occur for the most pleasant rated stimulus (high output AM on the palm), thus, high output AM is suggested as the most appropriate stimulus for applications that target all age groups, such as, haptic displays in cars, aviation haptic controls (73) and virtual barriers, such as a train platform warning system (70). Furthermore, concerning the therapeutics industry, where an ultrasound alternative to social touch would benefit the elderly in particular (70), the findings highlight that more research is needed to identify a C-LTM-optimal ultrasound stimulus, before testing for therapeutic benefits.

Somatosensory and affective neuroscience would benefit from further understanding of which mechanoreceptors and nerves are being activated in response to ultrasound stimuli. Microneurography investigation would, crucially, facilitate this objective with particular focus on FAI and FAII nerves in the median and lateral antebrachial nerves. In terms of research relating to the social touch hypothesis, it would be of importance to ascertain whether a C-LTM-optimal stimulus can be generated with ultrasound, using an objective method such as microneurography. This would include investigating the thermodynamic properties of ultrasound stimuli and whether the effect of airflow over the skin generates localised cooling via wind-chill effects. The field of haptics would benefit from further investigation into which stimuli are perceived as equal intensity as a function of peak ARP, time-averaged ARP, modulation and skin region. Furthermore, research concerning the nuances of intensity and pleasantness ratings across different age groups, (including children and adolescents) is recommended, in order to produce perceptually correct sensations between users. As a wider point of interest, it may be beneficial to the development of VR to investigate whether the Stratos Inspire device is capable of inducing the body ownership phenomenon documented in the rubber hand illusion (74). This seems a logical progression as the Stratos Inspire device has VR capabilities to produce an interactive environment, where the virtual hand is able to be stroked while producing the appropriate sensation on the user’s real hand (6).

## Conclusion

Intensity and pleasantness ratings of ultrasound stimulation were changed as a function of peak ARP, modulation, skin region and age; only intensity of AM stimulus was different between skin types; and the TEAQ only correlated with the pleasantness rating of stimulus four (C-LTM-optimal stimulus). The findings highlighted potential misinterpretation of force applied to the skin, which could further explain lower detection threshold of STM compared to AM (63) (66). While a range of stimulus intensities and pleasantness ratings were produced, the Stratos Inspire is also capable of creating stimuli on different skin types with almost equal sensation. These findings have implications for ultrasonic haptics, somatosensory affective research and virtual reality. The future study of haptic research will require multidisciplinary cooperation bringing together cognitive neuroscience and affective computing (75).

## References

1. McGlone F, Reilly D. The cutaneous sensory system. Neuroscience & Biobehavioral Reviews. 2010;34(2):148–59.

2. Le Pichon CE, Chesler AT. The functional and anatomical dissection of somatosensory subpopulations using mouse genetics. Frontiers in neuroanatomy. 2014;8:21.

3. Greenspan JD, LaMotte RH. Cutaneous mechanoreceptors of the hand: experimental studies and their implications for clinical testing of tactile sensation. Journal of Hand Therapy. 1993;6(2):75–82.

4. Kakuda N. Conduction velocity of low-threshold mechanoreceptive afferent fibers in the glabrous and hairy skin of human hands measured with microneurography and spike-triggered averaging. Neuroscience research. 1992;15(3):179–88.

5. Lumpkin EA, Caterina MJ. Mechanisms of sensory transduction in the skin. Nature. 2007;445(7130):858–65.

6. Ultraleap. Stratos Inspire. Available from: https://www.ultraleap.com/product/stratos-inspire/ [Accessed 05/12/19.]

7. Zotterman Y. Touch, pain and tickling: an electro-physiological investigation on cutaneous sensory nerves. The Journal of physiology. 1939;95(1):1–28.

8. Burgess P, Petit D, Warren RM. Receptor types in cat hairy skin supplied by myelinated fibers. Journal of neurophysiology. 1968;31(6):833–48.

9. Jänig W, Schmidt R, Zimmermann M. Single unit responses and the total afferent outflow from the cat’s foot pad upon mechanical stimulation. Experimental brain research. 1968;6(2):100–15.

10. Vallbo A, Johansson R. The tactile sensory innervation of the glabrous skin of the human hand. Active touch. 1978;2954:29–54.

11. Torebjörk H, Vallbo Å, Ochoa J. Intraneural microstimulation in man: Its relation to specificity of tactile sensations. Brain. 1987;110(6):1509–29.

12. Johansson RS, Vallbo A. Tactile sensibility in the human hand: relative and absolute densities of four types of mechanoreceptive units in glabrous skin. The Journal of physiology. 1979;286(1):283–300.

13. Knibestöl M, Vallbo ÅB. Single unit analysis of mechanoreceptor activity from the human glabrous skin. Acta Physiologica Scandinavica. 1970;80(2):178–95.

14. Hatzfeld C. Haptics as an interaction modality: Springer; 2014. 29–100.

15. Hao J, Bonnet C, Amsalem M, Ruel J, Delmas P. Transduction and encoding sensory information by skin mechanoreceptors. Pflügers Archiv-European Journal of Physiology. 2015;467(1):109–19.

16. Bolanowski SJ, Gescheider GA, Verrillo RT, Checkosky CM. Four channels mediate the mechanical aspects of touch. The Journal of the Acoustical society of America. 1988;84(5):1680–94.

17. Wiklund Fernström K. Physiological properties of unmyelinated low-threshold tactile (CT) afferents in the human hairy skin 2004.

18. Myers MI, Peltier AC, Li J. Evaluating dermal myelinated nerve fibers in skin biopsy. Muscle & nerve. 2013;47(1):1–11.

19. Johansson R, Vallbo Å. Spatial properties of the population of mechanoreceptive units in the glabrous skin of the human hand. Brain research. 1980;184(2):353–66.

20. Johansson RS, Vallbo ÅB. Tactile sensory coding in the glabrous skin of the human hand. Trends in neurosciences. 1983;6:27–32.

21. Woodbury CJ, Ritter AM, Koerber HR. Central anatomy of individual rapidly adapting low-threshold mechanoreceptors innervating the “hairy” skin of newborn mice: early maturation of hair follicle afferents. Journal of Comparative Neurology. 2001;436(3):304–23.

22. Johnson KO. The roles and functions of cutaneous mechanoreceptors. Current opinion in neurobiology. 2001;11(4):455–61.

23. Johnson KO, Yoshioka T, Vega–Bermudez F. Tactile functions of mechanoreceptive afferents innervating the hand. Journal of Clinical Neurophysiology. 2000;17(6):539–58.

24. Obrist M, Seah SA, Subramanian S, editors. Talking about tactile experiences. Proceedings of the SIGCHI Conference on Human Factors in Computing Systems; 2013.

25. Scheibert J, Leurent S, Prevost A, Debrégeas G. The role of fingerprints in the coding of tactile information probed with a biomimetic sensor. Science. 2009;323(5920):1503–6.

26. Verrillo RT. Age related changes in the sensitivity to vibration. Journal of gerontology. 1980;35(2):185–93.

27. Seah SA, Griffin MJ. Normal values for thermotactile and vibrotactile thresholds in males and females. International archives of occupational and environmental health. 2008;81(5):535–43.

28. Johansson RS. Tactile sensibility in the human hand: receptive field characteristics of mechanoreceptive units in the glabrous skin area. The Journal of physiology. 1978;281(1):101–25.

29. Verrillo RT. Investigation of some parameters of the cutaneous threshold for vibration. The Journal of the Acoustical Society of America. 1962;34(11):1768–73.

30. Olausson H, Lamarre Y, Backlund H, Morin C, Wallin B, Starck G, et al. Unmyelinated tactile afferents signal touch and project to insular cortex. Nature neuroscience. 2002;5(9):900.

31. Checkosky C, Bolanowski Jr S. Identifying possible neural codes for vibrotaction using the theory of temporal summation. The Journal of the Acoustical Society of America. 1989;85(S1):S63–S4.

32. Ribot-Ciscar E, Vedel J, Roll J. Vibration sensitivity of slowly and rapidly adapting cutaneous mechanoreceptors in the human foot and leg. Neuroscience letters. 1989;104(1-2):130–5.

33. Nordin M. Low-threshold mechanoreceptive and nociceptive units with unmyelinated (C) fibres in the human supraorbital nerve. The Journal of Physiology. 1990;426(1):229–40.

34. Wessberg J, Norrsell U. A system of unmyelinated afferents for innocuous mechanoreception in the human skin. Brain research. 1993;628(1-2):301–4.

35. Iriuchijima J, Zotterman Y. The specificity of afferent cutaneous C fibres in mammals. Acta Physiologica Scandinavica. 1960;49(2-3):267–78.

36. Vallbo Å, Olausson H, Wessberg J. Unmyelinated afferents constitute a second system coding tactile stimuli of the human hairy skin. Journal of neurophysiology. 1999;81(6):2753–63.

37. Wessberg J, Olausson H, Fernström KW, Vallbo ÅB. Receptive field properties of unmyelinated tactile afferents in the human skin. Journal of neurophysiology. 2003;89(3):1567–75.

38. Ackerley R, Wasling HB, Liljencrantz J, Olausson H, Johnson RD, Wessberg J. Human C-tactile afferents are tuned to the temperature of a skin-stroking caress. Journal of Neuroscience. 2014;34(8):2879–83.

39. Iggo A. Cutaneous mechanoreceptors with afferent C fibres. The Journal of physiology. 1960;152(2):337–53.

40. Löken LS, Wessberg J, McGlone F, Olausson H. Coding of pleasant touch by unmyelinated afferents in humans. Nature neuroscience. 2009;12(5):547.

41. Essick GK, James A, McGlone FP. Psychophysical assessment of the affective components of non-painful touch. Neuroreport. 1999;10(10):2083–7.

42. Pawling R, Trotter PD, McGlone FP, Walker SC. A positive touch: C-tactile afferent targeted skin stimulation carries an appetitive motivational value. Biological Psychology. 2017;129:186–94.

43. Ackerley R, Carlsson I, Wester H, Olausson H, Backlund Wasling H. Touch perceptions across skin sites: differences between sensitivity, direction discrimination and pleasantness. Frontiers in behavioral neuroscience. 2014;8:54.

44. Miles R, Cowan F, Glover V, Stevenson J, Modi N. A controlled trial of skin-to-skin contact in extremely preterm infants. Early human development. 2006;82(7):447–55.

45. McMullan S, Fisher L. Developmental progress of Romanian orphanage children in Canada. Canadian Psychology. 1992;33(2):504.

46. Ardiel EL, Rankin CH. The importance of touch in development. Paediatrics & child health. 2010;15(3):153–6.

47. Field T, Soliday B, Lasko D, Gonzalez N, Valdeon C. Touching in infant, toddler, and preschool nurseries. Early Child Development and Care. 1994;98(1):113–20.

48. Field T, Ironson G, Scafidi F, Nawrocki T, Goncalves A, Burman I, et al. Massage therapy reduces anxiety and enhances EEG pattern of alertness and math computations. International journal of neuroscience. 1996;86(3-4):197–205.

49. Pawling R, Cannon PR, McGlone FP, Walker SC. C-tactile afferent stimulating touch carries a positive affective value. PloS one. 2017;12(3).

50. Whitcher SJ, Fisher JD. Multidimensional reaction to therapeutic touch in a hospital setting. Journal of Personality and Social Psychology. 1979;37(1):87.

51. McGlone F, Wessberg J, Olausson H. Discriminative and affective touch: sensing and feeling. Neuron. 2014;82(4):737–55.

52. Green L. The trouble with touch? New insights and observations on touch for social work and social care. British Journal of Social Work. 2017;47(3):773–92.

53. Kyung K-U, Ahn M, Kwon D-S, Srinivasan MA, editors. A compact broadband tactile display and its effectiveness in the display of tactile form. First Joint Eurohaptics Conference and Symposium on Haptic Interfaces for Virtual Environment and Teleoperator Systems World Haptics Conference; 2005: IEEE.

54. Monnoyer J, Diaz E, Bourdin C, Wiertlewski M, editors. Ultrasonic friction modulation while pressing induces a tactile feedback. International Conference on Human Haptic Sensing and Touch Enabled Computer Applications; 2016: Springer.

55. Huisman G, Frederiks AD, van Erp JB, Heylen DK, editors. Simulating affective touch: Using a vibrotactile array to generate pleasant stroking sensations. International Conference on Human Haptic Sensing and Touch Enabled Computer Applications; 2016: Springer.

56. Nagi SS, Mahns DA. Mechanical allodynia in human glabrous skin mediated by low-threshold cutaneous mechanoreceptors with unmyelinated fibres. Experimental brain research. 2013;231(2):139–51.

57. Gursul D, Goksan S, Hartley C, Mellado GS, Moultrie F, Hoskin A, et al. Stroking modulates noxious-evoked brain activity in human infants. Current Biology. 2018;28(24):1380–1.

58. Culbertson H, Schorr SB, Okamura AM. Haptics: The present and future of artificial touch sensation. Annual Review of Control, Robotics, and Autonomous Systems. 2018;1:385–409.

59. Long B, Seah SA, Carter T, Subramanian S. Rendering volumetric haptic shapes in mid-air using ultrasound. ACM Transactions on Graphics (TOG). 2014;33(6):1–10.

60. Chilles J, Frier W, Abdouni A, Giordano M, Georgiou O, editors. Laser Doppler Vibrometry and FEM Simulations of Ultrasonic Mid-Air Haptics. 2019 IEEE World Haptics Conference (WHC); 2019: IEEE.

61. Ito M, Wakuda D, Inoue S, Makino Y, Shinoda H, editors. High spatial resolution midair tactile display using 70 kHz ultrasound. International Conference on Human Haptic Sensing and Touch Enabled Computer Applications; 2016: Springer.

62. Frier W, Ablart D, Chilles J, Long B, Giordano M, Obrist M, et al., editors. Using spatiotemporal modulation to draw tactile patterns in mid-air. International Conference on Human Haptic Sensing and Touch Enabled Computer Applications; 2018: Springer.

63. Brisben A, Hsiao S, Johnson K. Detection of vibration transmitted through an object grasped in the hand. Journal of neurophysiology. 1999;81(4):1548–58.

64. Raza A, Hassan W, Ogay T, Hwang I, Jeon S. Perceptually Correct Haptic Rendering in Mid-Air using Ultrasound Phased Array. IEEE Transactions on Industrial Electronics. 2019;67(1):736–45.

65. Fan L, Song A, Zhang H. Haptic Interface Device Using Cable Tension Based on Ultrasonic Phased Array. IEEE Access. 2020;8:880–91.

66. Takahashi R, Hasegawa K, Shinoda H, editors. Lateral modulation of midair ultrasound focus for intensified vibrotactile stimuli. International Conference on Human Haptic Sensing and Touch Enabled Computer Applications; 2018: Springer.

67. Rakkolainen I, Freeman E, Sand A, Raisamo R, Brewster S. A survey of mid-air ultrasound haptics and its applications. IEEE Transactions on Haptics. 2020;14(1):2–19.

68. Durkin J, Jackson D, Usher K. Touch in times of COVID-19: Touch Hunger hurts. Journal of Clinical Nursing. 2020.

69. Georgiou O, Limerick H, Corenthy L, Perry M, Maksymenko M, Frish S, et al. Mid-Air Haptic Interfaces for Interactive Digital Signage and Kiosks. Conference on Human Factors in Computing Systems; Glasgow: Association for Computing Machinery; 2019.

70. Mizutani S, Fujiwara M, Makino Y, Shinoda H, editors. Thresholds of Haptic and Auditory Perception in Midair Facial Stimulation. 2019 IEEE International Symposium on Haptic, Audio and Visual Environments and Games (HAVE); 2019: IEEE.

71. Osczevski RJ. The basis of wind chill. Arctic. 1995:372–82.

72. Beattie D, Georgiou O, Harwood A, Clark R, Long B, Carter T. Mid-Air Haptic Textures from Graphics. USA. 16/734, 479 (Patent). 2019.

73. Girdler A, Georgiou O. Mid-Air Haptics in Aviation--creating the sensation of touch where there is nothing but thin air. arXiv [preprint]. 2020.

74. Horiuchi Y, Yoshida K, Makino Y, Shinoda H, editors. Rubber hand illusion using invisible tactile stimulus. 2017 IEEE World Haptics Conference (WHC); 2017: IEEE.

75. Eid MA, Al Osman H. Affective haptics: Current research and future directions. IEEE Access. 2015;4:26–40.

